# HaploCatcher: An R Package for Prediction of Haplotypes

**DOI:** 10.1101/2023.07.20.549744

**Authors:** Zachary James Winn, Emily Hudson-Arns, Mikayla Hammers, Noah DeWitt, Jeanette Lyerly, Guihua Bai, Paul St. Amand, Punya Nachappa, Scott Haley, Richard Esten Mason

## Abstract

Wheat (*Triticum aestivum* L.) is crucial to global food security, but is often threatened by diseases, pests, and environmental stresses. Wheat stem sawfly (*Cephus cinctus* Norton) poeses a major threat to food security in the United States, and solid-stem varieties, which carry the stem-solidness locus (*Sst1*), are the main source of genetic resistance against sawfly. Marker-assisted selection uses molecular markers to identify lines possessing beneficial haplotypes, like that of the *Sst1* locus. In this study, an R package titled "HaploCatcher" was developed to predict specific haplotypes of interest in genome-wide genotyped lines. A training population of 1,056 lines genotyped for the *Sst1* locus, known to confer stem solidness, and genome-wide markers was curated to make predictions of the *Sst1* haplotypes for 292 lines from the Colorado State University wheat breeding program. Predicted *Sst1* haplotypes were compared to marker derived haplotypes. Our results indicated that the training set was substantially predictive, with kappa scores of 0.83 for k-nearest neighbors and 0.88 for random forest models. Forward validation on newly developed breeding lines demonstrated that a random forest model, trained on the total available training data, had comparable accuracy between forward and cross-validation. Estimated group means of lines classified by haplotypes from PCR-derived markers and predictive modeling did not significantly differ. The HaploCatcher package is freely available and may be utilized by breeding programs, using their own training populations, to predict haplotypes for whole genome sequenced early generation material.

**CORE IDEAS:**

1. Identification, introgression, and frequency increase of large effect loci are important for cultivar development.
2. The *Sst1* locus has a significant effect on cutting score in fields exposed to sawfly infestation.
3. Historical genetic information can be utilized to predict haplotypes for lines which have genome-wide genetic data.
4. An R package, HaploCatcher, has been developed to facilitate this analysis in other programs.

## INTRODUCTION

Common bread wheat (*Triticum aestivum* L.) consumption represents nearly 20% of human caloric intake; however, current genetic gain of wheat grain yield is insufficient to meet the rise in demand as the global population increases (Poole et al., 2021; Ray et al., 2013; Shiferaw et al., 2013). Two major threats to grain yield stability in wheat are diseases and pests. One such pest, which presents a major risk to yield stability in the United States northern Great Plains and Mountain West regions, is wheat stem sawfly (*Cephus cinctus* Norton). In terms of domestic losses, the Colorado winter wheat growing region lost approximately 32.7 and 31.2 million dollars’ worth of wheat in the years 2020 and 2021, respectfully (Erika et al., 2023). Yield losses for infested hollow-stem varieties can be anywhere from 90-120 kg ha^−1^ for spring wheats, potentially resulting in multi-million dollar losses annually (Beres et al., 2007). Furthermore, Beres et al (2011) estimated that sawfly infestation may cost 350 million dollars annually to the United States northern Great Plains and Canadian provinces, making it a major concern of consumers and producers alike.

The wheat stem sawfly is an insect native to North America that infests wheat by ovipositing eggs within the stem of wheat plants from late May to early June (Weiss & Morrill, 1992). Once the egg has been deposited into the stem, the larva emerges and feeds upon the parenchyma and vascular tissue inside the stem (Weiss & Morrill, 1992). After receiving the correct combination of physical and photoperiodic signals (Holmes, 1975), the larvae will migrate downward from its hatching site to an area of the stem near the soil surface and create a notch, which is known as a hibernaculum, that it fills with the excrement of digested plant material (frass) (Weiss & Morrill, 1992). The wheat stem tends to break at the notch development site, causing the substantial lodging of affected plants that gives the pest its common name. The larvae will enter a period of diapause to then pupate and emerge from the hibernaculum in the following year (Beres et al., 2011).

Although wheat stem sawfly are weak fliers which tend to oviposit near the area of emergence (Beres et al., 2011; Weiss & Morrill, 1992), their distribution is wide and integrated pest management is challenging. Removal of stubs was once recommended as a key cultural control of wheat stem sawfly (Fletcher, 1904), but contemporary research proved that that method was ineffective (Beres et al., 2011). Rotations of wheat followed by fallows also appeared to increase infestation rates, so it has been suggested to use a non-host in rotation as a “trap crop” (Beres et al., 2011; Seamens, 1929). More recently, trap crops have been suggested as a management tool where trap crops are planted as a border around hollow-stem varieties to act as a buffer-zone and prevent infestation of higher yielding hollow-stem varieties (Beres et al., 2009; Peirce, Cockrell, Ode, et al., 2022).

Solid-stem varieties of wheat have been available since the mid-twentieth century (Peirce, Cockrell, Mason, et al., 2022; Weiss & Morrill, 1992). In solid-stem varieties, undifferentiated parenchyma cells create a solid pith within the stem (Berzonsky et al., 2003) and this lessens the severity of yield losses in wheat plants (Beres et al., 2007). The genetic architecture of stem solidness appears oligogenic, with large effect loci being the main contributors to solidness (Peirce, Cockrell, Mason, et al., 2022). One gene found responsible for solidness is *Sst1* (Nilsen et al., 2020, 2017), which was first identified in a QTL study conducted by Cook et al (2004) on the long arm of wheat chromosome 3B (*Qss.msub-3BL*). Stem solidness is thus caused, in part, by tandem repeats of the *TdDof* gene coding sequence which lead to the filling in of the pith within the wheat stem (Nilsen et al., 2020).

While visual rating of stem solidness can be a reliable method for selecting lines that express solid-stem phenotypes, many wheat breeders in the United States utilize molecular markers to haplotype the region containing *Sst1* in a process termed marker-assisted selection. More recently, the United States Department of Agriculture (USDA) Central Small Grains Genotyping Lab located in Manhattan, Kansas has been producing haplotype information on many large effect loci, including *Sst1*, for the lines entered into the Southern Regional Performance Nursery (SRPN) and Regional Germplasm Observation Nursery (RGON) [https://www.ars.usda.gov/plains-area/lincoln-ne/wheat-sorghum-and-forage-research/docs/hard-winter-wheat-regional-nursery-program/research/]. This service performed by the USDA lab is conducted to assist breeders in releasing lines with the solid-stem trait, and, as a result, it has also created a backlog of information on lines in the SRPN and RGON lines with known *Sst1* haplotypes.

Moreover, the lines in the SRPN and RGON have been characterized for genome-wide single nucleotide polymorphisms (SNPs) by the Colorado State University (CSU) Wheat Breeding Program on an annual basis for more than a decade. Winn et al (2022) described a method where historical molecular and haplotype data are utilized to produce accurate haplotype predictions on lines which only have genome-wide molecular data by characterizing either homozygous resistant or homozygous susceptible varieties

In the current work we sought to (1) produce a deployable R statistical software compatible package to perform an analysis similar to the one performed in Winn et al (2022), (2) demonstrate that the analysis preforms similarly in an unrelated germplasm pool for a different locus than those explored in Winn et al (2022), (3) predict the *Sst1* haplotypes of genome-wide genotyped individuals, and (4) compare the effect of predicted *Sst1* verses genotyped *Sst1* on wheat stem sawfly related phenotypes in Colorado State University hard winter wheat germplasm.

## MATERIALS AND METHODS

### Germplasm

Two separate sets of germplasm were utilized in this study. The first population used in this study was a historical panel of lines submitted to the SRPN and RGON. This panel of lines consisted of 1,056 distinct genotypes, and all lines in the panel were genotyped genome-wide for SNPs and haplotyped via a diverse panel of markers for the *Sst1*/*Qss.msub-3BL* locus. The second population utilized in this study represented contemporary lines in the CSU Wheat Breeding Program from the 2022 advanced yield nursery (AYN) and the 2022 wheat stem sawfly solid stem panel (WSS), which were phenotyped for sawfly reaction traits, genotyped for SNPs across the genome, and screened with kompetative allele specific polymerase chain reaction (KASP) assays for the *Sst1* locus. The AYN consisted of 107 distinct genotypes and the WSS consisted of 185 distinct genotypes (292 total genotypes). Individuals in the WSS had not gone through any phenotypic or marker-assisted selection for solid stem or wheat stem sawfly resistance, while individuals in the AYN had already undergone one generation of field selection for resistance.

### Phenotyping

In the 2021-2022 wheat growing season, the AYN and WSS were planted in Akron, Colorado and a second location of the AYN was planted in New Raymer, Colorado. These sites were selected for sawfly trials due to the historical presence of sawfly within these regions and the consistent infestation that they receive (Cockrell et al., 2021; Irell & Peairs, 2014). Furthermore, in areas surrounding the field sites, 100 sweeps were taken along the field edge bordering an adjacent wheat stubble field. Sampling began during mid-jointing and continued weekly until no adult sawfly were found in sweep samples. This data further confirms if infestation pressure was adequate for data collection (Nachappa, 2023; Nachappa & Peirce, 2022).

In each site, all plots were sown in mid-September using a 1.5m wide no-till drill seeder that was guided by a cable and had 4.9m spacing between centers. Following spring green up, centers were pruned using glyphosate (Bayer, St Louis, Missouri, USA) applied by a 1.2m wide hooded sprayer. After end trimming, this resulted in a measurable area of 1.5m by 3.7m. The AYN and WSS were planted in partially replicated designs arranged in rows and columns, with repeated checks included at both row and column levels.

After physiological maturity, when lodging due to sawfly cutting was apparent, a visual cutting score was assigned to each plot in each location. The visual cutting score was assigned as an index of percent plot affected by cutting, which is the physical process by which insect injury detaches most of the wheat stem from the base of the plant. Visual scores were assigned via an ordinal scale ranging from 1-9, where one is fully resistant and erect despite wheat stem sawfly pressure, and nine indicates the whole plot is affected, cut, and prostrate.

### Genome-Wide Genotyping

Ten seeds were planted for each line and a 2-3 cm of leaf tissue sample was taken from each plant and bulked for DNA extraction. Genomic DNA was extracted from the samples using MagMax (ThermoFisher Scientific; Waltham, Massachusetts, USA) plant DNA kits following the manufacturer’s instructions and quantified using PicoGreen (ThermoFisher Scientific; Waltham, Massachusetts, USA) kits. Extracted DNA was normalized to a concentration of 20 ng µL^−1^ and sequencing libraries were prepared following the protocol established by Poland et al (2012). The multiplexed libraries were sequenced on a NovaSeq 6000 (Illumina, San Diego, California, USA) sequencer at 384-plex density per lane. The resulting reads were aligned to the International Wheat Genome Sequencing Consortium (IWGSC) wheat reference sequence RefSeq v2.0 (Appels et al., 2018) using the burrow-wheeler aligner (Li & Durbin, 2009).

The TASSEL 2.0 standalone pipeline (Glaubitz et al., 2014) was used to process the reads obtained from alignment, and markers were organized into compressed variant calling format files (Danecek et al., 2011). Initial variant calling format files were filtered using the following parameters: monomorphic SNPs, insertions, and deletions were removed, SNPs with 85% or less missing data were retained, SNPs with a read depth of more than one or less than 100 were retained, SNPs with a minimum allele frequency of less than 5% were removed, SNPs with more than 10% heterozygosity were removed, and all unaligned SNPs were removed. After filtration, missing data were imputed using the Beagle algorithm V5.4 (Browning et al., 2018), and a synthetic wheat biparental cross between ’W7984’ and ’Opata’ was used to derive a recombination distance-based map for imputation (Gutierrez-Gonzalez et al., 2019).

### Historical Haplotype Information Curation

Information on the *Sst1* locus was curated from historical marker calling files generated by the USDA Central Small Grains Genotyping Lab [https://www.ars.usda.gov/plains-area/lincoln-ne/wheat-sorghum-and-forage-research/docs/hard-winter-wheat-regional-nursery-program/research/]. The information for lines in both the RGON and SRPN was standardized to a biallelic haplotype of homozygous *Sst1*, heterozygous, and homozygous wildtype represented as “+/+”, “+/-”, and “-/-”, respectively.

Haplotype calls within the CSU wheat breeding program were made using a single marker identified as diagnostic for the *Sst1* locus. Extracted and purified DNA, ranging between 50n and 150 ng uL^−1^ were plated in 96-well plates in 4 uL volumes. Plates contained both test genotype DNA as well as positive, heterozygous, negative, and non-template controls. Each well 4 uL of 2X KASP (LGC Genomics, Middlesex, UK) reaction mixture and 0.11 uL of KASP primer assay mixture. The assay mixture contained an equal mixture of 100 uM of FAM and HEX fluorescence labeled forward primers, as well as 2.5 concentration of 100 uM reverse primer, suspended in molecular grade sterile water (Table 1). Assays were run on a Bio-Rad (Bio-Rad; Hercules, California, USA) CFX96 RT PCR machine and results were read using a single endpoint measurement of florescence. Haplotype calls were made by visual discrimination of florescence groupings. The frequency of allelic states for *Sst1* in the training and testing set are also provided (Table 2).

**Table 1.**
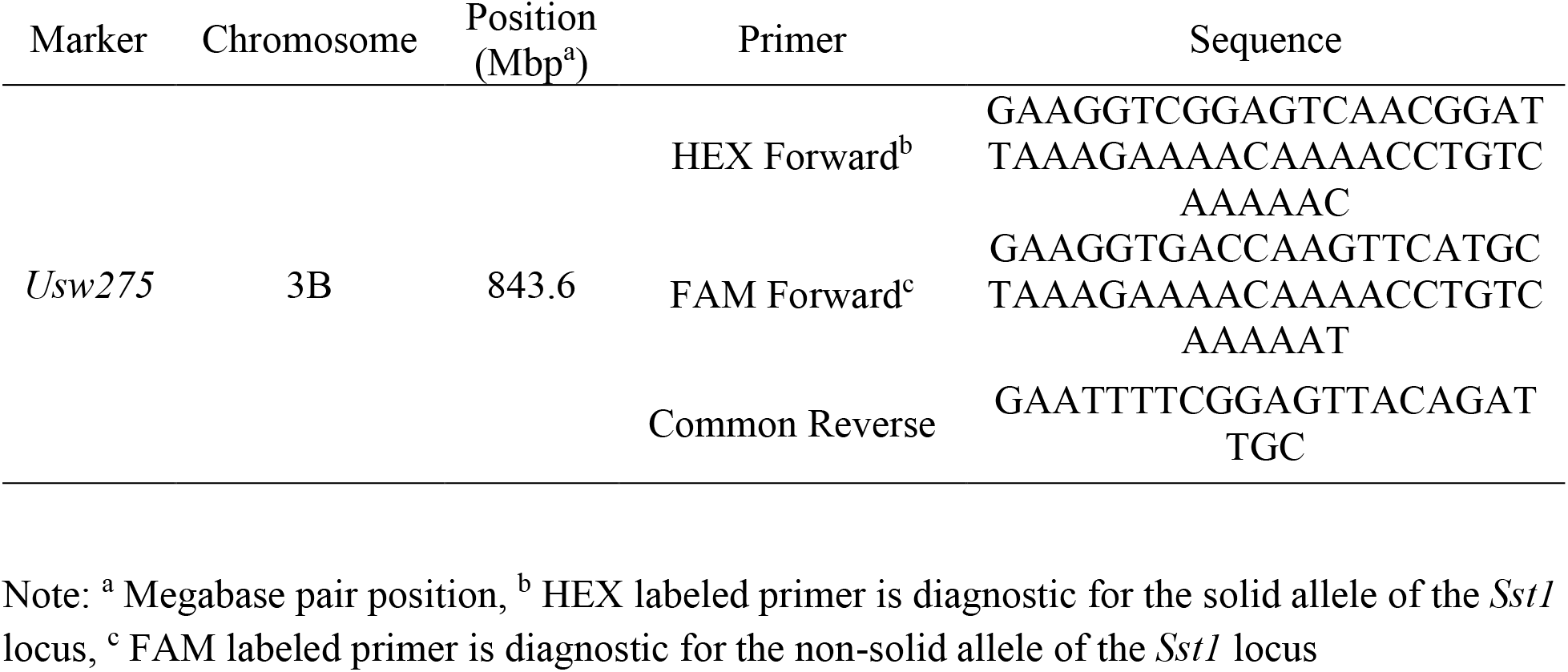
List of primers and sequences for the *Usw275* marker.

**Table 2.** Number of observations and frequency of *Sst1* haplotypes in the training and test population.

| Haplotype | Training |  | Testing |  |
| --- | --- | --- | --- | --- |
|  | n<sup>a</sup> | Frequency | n | Frequency |
| +/+<sup>b</sup> | 303 | 0.29 | 156 | 0.53 |
| +/-<sup>c</sup> | 104 | 0.10 | 25 | 0.09 |
| -/-<sup>d</sup> | 649 | 0.61 | 111 | 0.38 |

Note: <sup>a</sup> number of observations, <sup>b</sup> homozygous *SstI* haplotype, <sup>c</sup> heterozygous *SstI* haplotype, <sup>d</sup> homozygous wildtype haplotype

### Package Development and Analysis Pipeline

The “HaploCatcher” package was developed using the “devtools” package (Wickham et al., 2022) in R statistical software (R Core Team, 2022) via the RStudio (Posit; Boston, Massachusetts, USA) development environment on a computer with a Microsoft® Windows operating system. Data inputs required of the package are a marker matrix containing both individuals in the training population and those in the testing population, a historical haplotype classification file for individuals in the training population, and a marker information file which denotes the name, chromosome, and position of each marker in the genotypic matrix (Figure 1A). The package is comprised of several core functions which are then streamlined into the function “auto_locus”. The “auto_locus” function conducts a similar analysis pipeline to Winn et al (2022) through the “caret” package (Kuhn, 2008, 2022), while requiring minimal intervention from users (Figure 1B).

**Figure 1.**
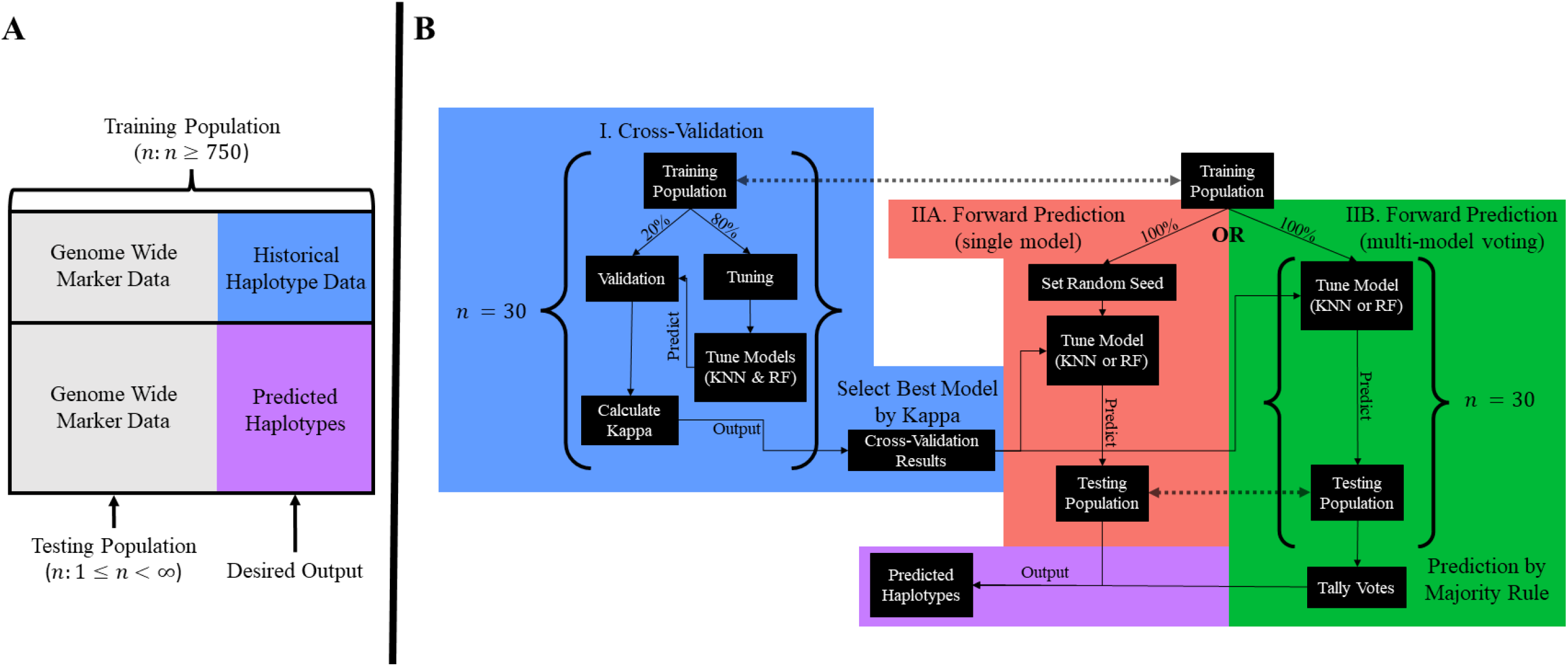
A diagram of [A] input data structure and [B] the “auto_locus” function pipeline. Panel [A] shows a total data set that is partitioned into a training and test population. The training population in panel [A] shows a population of individuals, that is suggested to be comprised of more than 750 individuals, which have both genome-wide marker and historical haplotype data. The testing population in panel [A] shows a testing population, which may be any size greater than zero, which only has genome-wide marker data. Panel [B] shows the workflow of the “auto_locus” function. In the cross-validation step [I], the total training population is split in a user defined way (default is 80:20 split) and the 80% tuning population is used to train and select optimal hyper-parameters for a k-nearest neighbors (KNN) and random forest (RF) model. The trained models are then used to predict the haplotype of the validation population. The predicted haplotype is then compared to the ‘true’ haplotype and kappa (and accuracy) are calculated. This is repeated a user set number of times (default is 30). The best performing model based on accuracy or kappa (default is kappa) is then taken as the model to be used in forward prediction. There are two options post cross-validation: [IIA] a single model with a set seed for repeatability or [IIB] a user set number of random models (default is 30) used to create a consensus haplotype prediction.

The “auto_locus” function is comprised of two major phases: cross validation by random partitioning into a user specified ratio (default is 80:20 training-testing; Figure1B - Step I) over a set number of permutations (default is 30) and forward prediction of training population candidates by the best model in cross validation (Figure1B-Step IIA and Figure1B- Step IIB). Prediction of training population haplotypes can be performed by either setting a random seed for reproducibility and performing the model once (Figure 1B - Step IIA) or by running the optimal model with no set seed over a user specified number of permutations (default is 30; Figure 1B - Step IIB) and producing a haplotype prediction by majority rule.

Cross-validation results were visualized using functions from the packages “ggplot2” and “patchwork” (Pedersen, 2022; Wickham et al., 2016). Both the cross-validation and forward prediction by voting steps in the “auto_locus” function can be run either sequentially or in parallel using the R packages “parallel”, “doParallel”, and “foreach” (Microsoft & Weston, 2022a, 2022b; R Core Team, 2022). Users can specify if the analysis is to be done in parallel (default argument is FALSE) or sequentially. Users may also define the number of processing cores desired for analysis or use the default setting which uses the function “detectCores” from the parallel package to determine the number of system cores and subtract that value by one.

The computer used for development of the package had a hexacore 2.6GHz Intel® (Intel; Santa Clara, California, USA) i7-10750H processor with 12 logical processors, 32 gigabytes of DDR4 RAM and a dedicated NVIDA (NVIDA; Santa Clara, California, USA) GeForce® RTX 2070 graphics card. Using the example datasets available in the HaploCatcher package, the “auto_locus” function performed in parallel with 100 permutations of cross-validations and 100 votes for majority rule resulted in a total runtime of eight minutes and 36 seconds.

### Statistical Analysis

All statistical analysis was conducted in R statistical software version 4.2.2 (R Core Team, 2022). Cutting visual score data was checked for normality by visualization of the distribution of observations using histograms and QQ-plots. Upon evaluation, all data exhibited near-normality or somewhat skewed normal distributions. Mixed linear models were run using the function “mmer” in the package “sommer” (Covarrubias-Pazaran, 2016, 2018). Across locations the following model was utilized to estimate the effect of the *Sst1* locus:

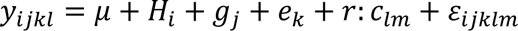

Where *y*_*ijklm*_ is the response, *μ* is the population mean, *H*_*i*_ is the haplotype fixed effect of the i^th^ haplotype, *g*_*j*_ is the genotypic random effect of the j^th^ genotype effect whose variance is defined by the additive relationship matrix among individuals derived by markers (VanRaden, 2008), *e*_*k*_ is the random environment effect of the k^th^ environment that is identically and independently distributed across levels, *r*: *c*_*lm*_ is the random row by column interaction effect of the l^th^ row and the j^th^ column whose variance is defined by the two-dimensional penalized tensor-product of spline relationship between row and column effects as described by Lee et al (2013), and *ε*_*ijklm*_ is the residual error that is identically and independently distributed across all levels.

To compare KASP genotyped haplotype vs machine learning predicted haplotype effects, the same mixed linear model was run twice to estimate an *H*_*i*_ haplotype fixed effect first using the “true” haplotype calls derived by KASP genotyping and then using the machine learning predicted haplotype information. Fixed effect group mean estimates for both the observed and predicted haplotype effects were estimated via the “predict.mmer” function in the package “sommer”. Visual comparison of effect estimates was summarized using functions from the “ggplot2” package.

Narrow-sense, per-plot, genomic heritability (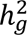) of cutting visual score ratings within environment were estimated using the following mixed linear model:

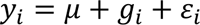

Where *y*_*i*_ is the observation, *μ* is the population mean, *g*_*i*_ is the random genotype effect of the i^th^ genotype whose variance is defined by the marker-derived additive relationship matrix calculated by the “A.mat” function from the “sommer” package, and ***ε***_**i**_ is the residual error whose variance is identically and independently distributed. Variance components were used in the function “vpredict” in the “sommer” package to estimate 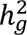 using the following formula:

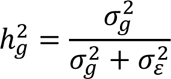

Where 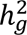 is the narrow-sense, per-plot, genomic heritability, 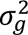 is the genotypic variance, and 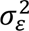 is the residual error variance.

Importance of defining variables (genome-wide SNP markers) in KNN and RF algorithms was calculated for each iteration of the 100 permutations of cross-validation by using the function “varImp” in the “caret” package. Variable importance, or more aptly put the importance of genome-wide SNP markers in defining haplotypes, was scaled between zero and 100 for comparability across models and the average importance of markers across all permutations was reported in images generated by the “ggplot2” package. Linkage disequilibrium (LD) was calculated for all markers identified as important using the function “LD” in the package “gaston” and results were reported in images derived by functions in the “ggplot2” package (Perdry & Dandine-Roulland, 2018).

Confusion matrices were calculated by comparing the predicted haplotype to the observed haplotype state in the WSS and AYN combined via the function “confusionMatrix” in the “caret” package. Model performance parameters were calculated across iterations of the 100 permutations of cross-validation and the forward prediction of the WSS and AYN. The reported measures of model performance were accuracy, sensitivity, specificity, and unadjusted Cohen’s kappa (McHugh, 2012). Accuracy was calculated as:

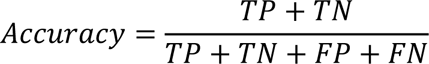

Where TP is the number of true positive cases, TN is the number of true negative cases, FP is the number of false positive cases, and FN is the number of false negative cases. Sensitivity and specificity were calculated as such:

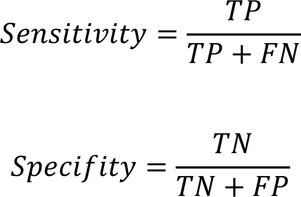

Cohen’s kappa was calculated as:

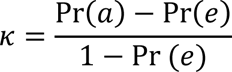

Where Pr(a) is the probability of observed agreement, and Pr(e) represents the expected rate of chance agreement. Kappa is often considered more robust than accuracy as a measurement parameter of reliability in categorization models because it is not easily biased by sample size (McHugh, 2012).

Kappa may be understood as a measurement which is bound between −1 and 1 where a value of 1 represents a perfectly categorizing model, 0 is the same as chance agreement, and a value of −1 is categorization that is worse than chance agreement (Viera et al., 2005). Historically, a kappa value of 0.8 to 1 is considered to be either “substantial” to “almost perfect” in its predictive ability (Landis & Koch, 1977). All model parameters were either reported in tables or visualized using functions from the “ggplot2” package.

## RESULTS

### *Sst1* Prediction Cross-Validation

Cross-validation indicated that the training data was well suited for analysis and substantially predictive based on reported kappa values (Figure 2). Over the 100 permutations of cross-validation, the average kappa value for the KNN model was *κ* = 0.83 and *κ* = 0.88 for RF. The average accuracy for the KNN model was 0.91 and 0.94 for RF. By-class sensitivity varied by haplotype. For homozygous *Sst1* calls, KNN had a mean sensitivity of 0.84 and RF had a mean sensitivity of 0.91. For heterozygous *Sst1* calls, KNN had a mean sensitivity of 0.82 and RF had a mean sensitivity of 0.81. For homozygous wildtype calls, both KNN and RF had a sensitivity of 0.96.

**Figure 2.**
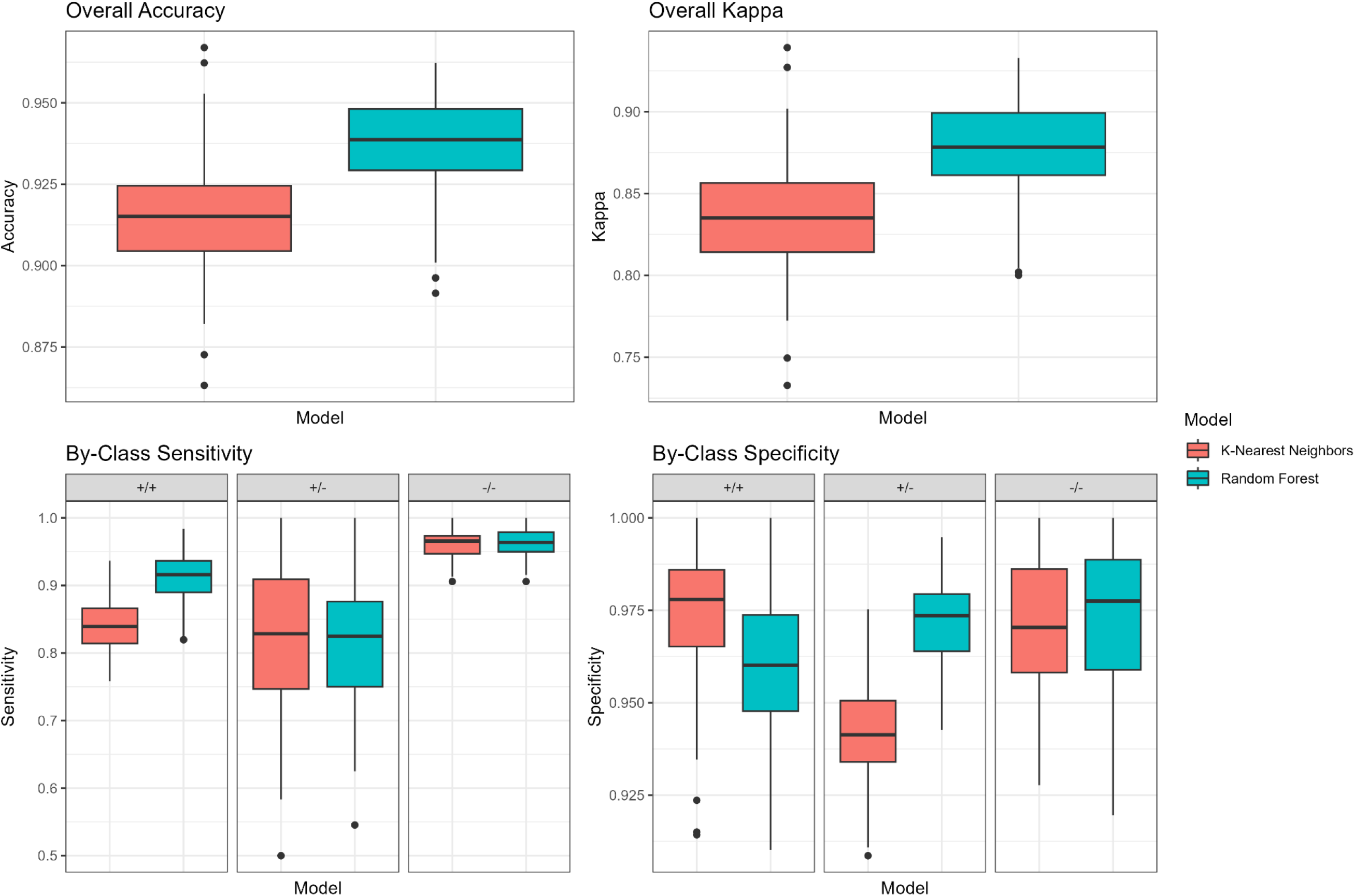
A visualization output by the “auto_locus” function of overall accuracy **(A)**, kappa, and by-class sensitivity and specificity value distributions over 100 permutations of cross validation. The figure legend on the right of the total figure displays the color that corresponds to which model. The top left panel displays the overall accuracy of each model in boxplots. The top right panel displays the overall kappa of each model in boxplots. The bottom left panel displays the by-class sensitivity values in boxplots for homozygous *Sst1* individuals (+/+), heterozygous individuals (+/-) and homozygous wildtype individuals (-/-). The bottom right panel displays the specificity of each model for each classification in boxplots. The x-axis is the model in each figure. The y-axis corresponds to the value of interest displayed within the graph.

Specificities had a narrow range among haplotype classifications and models. Average specificities for homozygous *Sst1* individuals were 0.97 for KNN and 0.96 for RF. Specificities for heterozygous *Sst1* individuals were 0.94 for KNN and 0.97 for RF. Specificities for homozygous wildtype individuals were 0.97 for both KNN and RF. These results indicate that KNN tended to under-identify true negatives in heterozygous cases, meaning that it tended to overclassify non-heterozygous individuals as heterozygous. Furthermore, the lower sensitivity scores of both the RF and KNN models (as compared to the higher sensitivity in the homozygous cases) indicates that both models were not as well suited for classifying heterozygous individuals as they were for homozygous individuals. Based on highest achieved average kappa value, the random forest model was selected for use in forward prediction.

Models in cross-validation mainly selected markers in or near the known region of *Sst1*, however there were two outliers on the distal short arm of 3B at approximately 34 megabase pairs (Mbp) and 159 Mbp (Figure 3B). When looking at LD among the markers selected for use in the models, it appears that the outlier markers and markers in the 828-852 Mbp region share minor-to-substantial LD (*r*^2^: 0.20 < *r*^2^ < 0.70; Figure 3A). More specifically, the LD appears to be very strong between these two outliers and markers in the 848-850 Mbp region (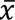 ≈ 0.66) which indicates that the markers may not be inherited independently. This may be the result of misalignment of markers to the wrong arm of the 3B chromosome. Alternatively, this may be a signature of true linkage disequilibrium, indicating that some region on the short arm of 3B is being inherited frequently with the *Sst1* locus.

**Figure 3.**
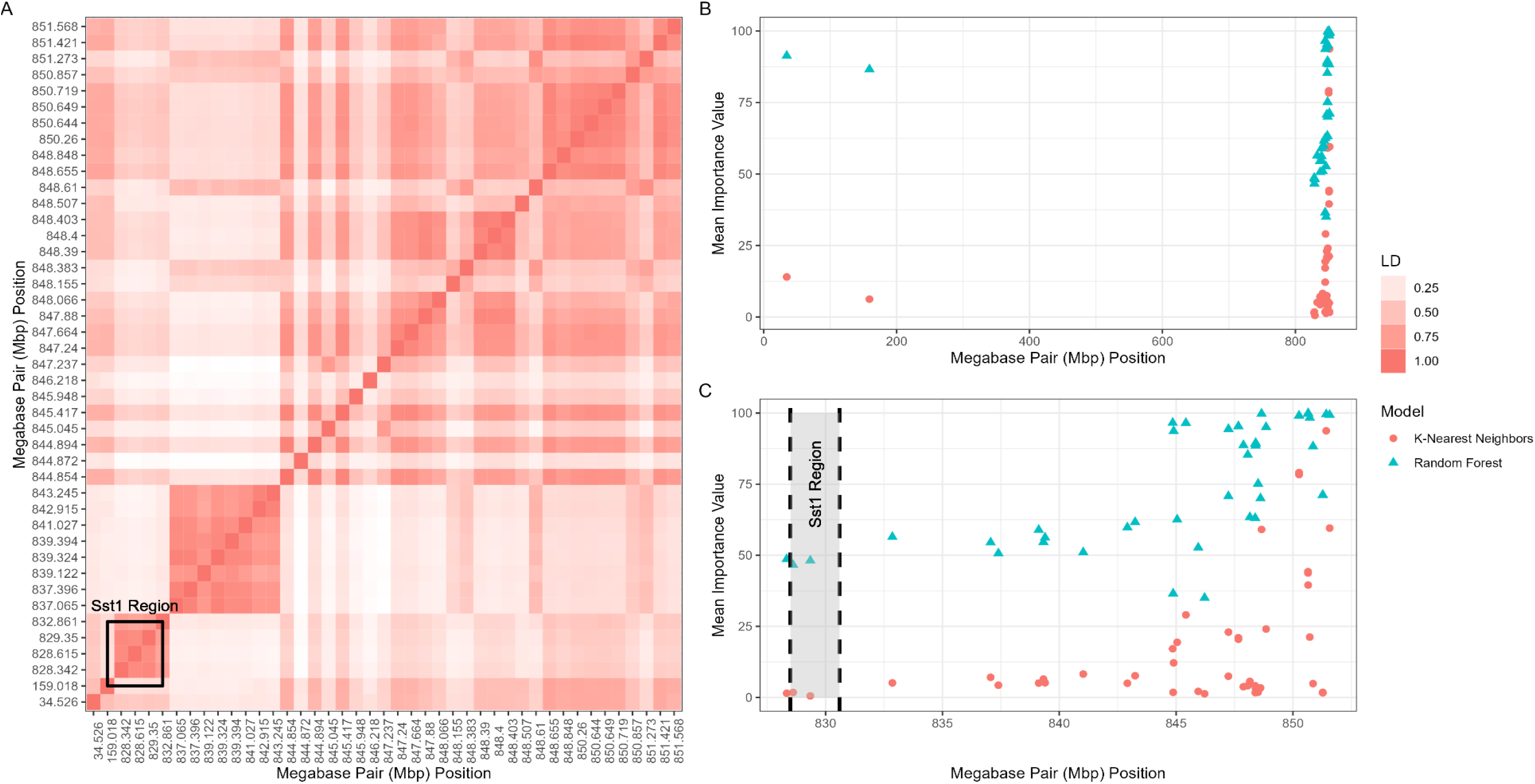
A visualization of (A) linkage disequilibrium (LD) among the most important markers identified between the k-nearest neighbors and random forest model, (B) the importance values of markers across the genome and (C) the importance values of markers proximal to the known position of *Sst1*. Panel (A) displays the linkage disequilibrium of each marker identified as important by the models. The x and y axes display the marker megabase pair (Mbp) position of each marker. The color within the plot on panel (A) that corresponds with the figure legend located on the right indicates the magnitude of LD between those markers. The known location of *Sst1* falls within the black box. The graph in panel (B) shows the importance of markers averaged over the 100 iterations. The y-axis displays the importance value of the marker which is represented by the colored dot. The x-axis represents the Mbp position of the marker. The point color corresponds with which model the point belongs to, which is denoted by the figure legend to the right. The graph in panel (C) is a zoomed in version of panel (B) where the known location of *Sst1* is labeled with a gray shaded box flanked by dashed lines.

When looking at derived importance values within the region, it appears that those outlier markers on the distal short arm of 3B are highly important (*x* > 0.75) for the RF model and less so for the KNN model (*x* < 0.25, Figure 3B). Taking a closer look at the known location of *Sst1*, it appears that markers within the region are moderately important (*x* ≈ 0.50) in RF models while they were non-important for KNN models (*x* < 0.10) (Figure 3B). Interestingly, the most important markers (*x* ≥ 0.95) identified by KNN and RF models were in the 845-853 Mbp region.

This region of highly important markers is located nearly 15-20 Mbps away from the known location of *Sst1*. However, historical markers used to haplotype the *Sst1* in the RGON and SRPN are not in perfect linkage with the causal polymorphism. Furthermore, the marker used by the CSU wheat breeding program, which is diagnostic of the *Sst1* locus, is found at approximately 843 Mbp, which is directly adjacent to the most important markers for classification. These results may be due to the use of haplotype designations derived from markers which do not lie within or in direct proximity to the *Sst1* locus. Regardless, model performance parameters, namely kappa, indicate that both models are capable of “substantial predictions” using historical scales for kappa interpretation (Landis & Koch, 1977).

### *Sst1* Prediction Forward Validation

Forward validation on the WSS and AYN using a RF model trained on the total available training data produced similar results to that of cross-validation (Figure 4). Accuracies for homozygous wildtype and *Sst1* individuals were 0.95 and 0.93, respectively. As observed in the cross-validation results, the accuracy for identification of heterozygous individuals was lower at 0.84. Specificities for homozygous *Sst1*, heterozygous *Sst1*, and homozygous wildtype were 0.96, 0.92, and 0.99. Sensitivities followed the same trend as cross-validation, where the true positive identification rate for homozygous *Sst1* and wildtype individuals was higher (0.95 and 0.85, respectively) than identification of heterozygous individuals (0.75).

**Figure 4.**
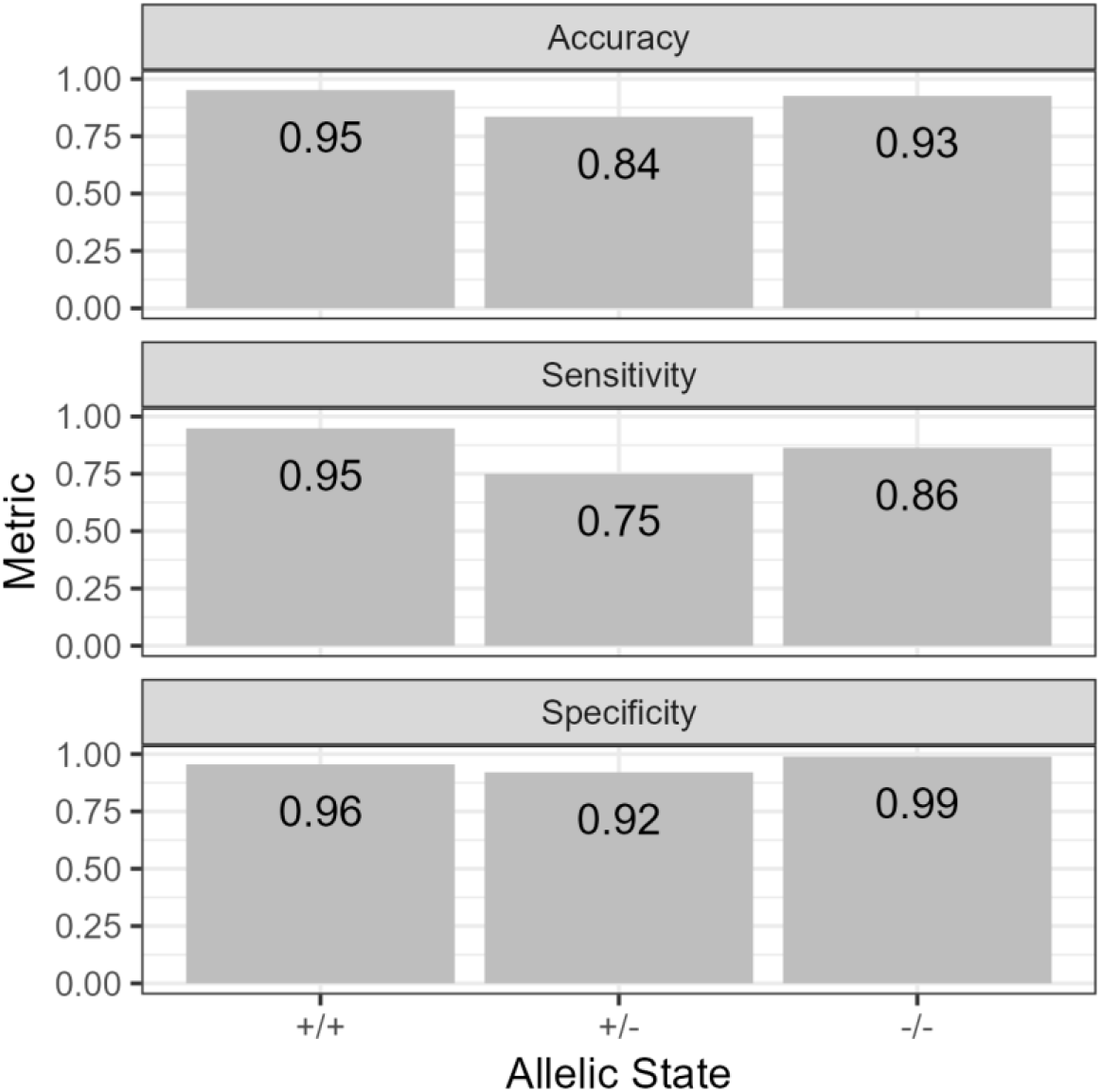
Visualization of performance parameters of predictions by a random forest model trained on all available training data. Each subgraph represents a separate measurement of model performance. The y-axis displays the magnitude of the metric displayed in each subgraph. The allelic state on the x-axis denotes individuals who are homozygous *Sst1* (+/+), heterozygous (+/-), and homozygous wildtype (-/-). The value of the metric for each allelic state is displayed within each bar.

Based on the confusion matrix (Table 3) of predicted vs observed haplotypes, the RF algorithm misidentified heterozygous individuals as wildtype frequently. Homozygous wildtype individuals were most often correctly identified (two cases misidentified), followed by homozygous *Sst1* individuals (seven cases misidentified). These results may indicate that, while not completely uninformative, these methods may be best suited for identifying homozygous individuals, like in Winn et al (2022), rather than trying to identify heterozygous individuals as well.

**Table 3.** Confusion matrix of predicted *Sst1* haplotypes calls vs haplotypes calls made by kompetative allele specific polymerase chain reaction (KASP)

|  |  | KASP (Observed) |  |  |
| --- | --- | --- | --- | --- |
|  |  | +/+ | +/- | -/- |
| Predicted | +/+<sup>a</sup> | 147 | 7 | 0 |
|  | +/-<sup>b</sup> | 6 | 19 | 15 |
|  | -/-<sup>c</sup> | 2 | 0 | 96 |

Note: <sup>a</sup> homozygous *SstI* calls, <sup>b</sup> heterozygous *SstI* calls, <sup>c</sup> homozygous wildtype calls

### Effect of *Sst1*: Predicted vs Genotyped Haplotype

Visual examination of distributions for both locations of the AYN revealed somewhat skewed data distributions while observations from the single location of the WSS followed an approximately normal distribution (Figure 5). Notably, the AYN exhibited a distribution skewed towards lower values of cutting visual scores at both Akron, CO and New Raymer, CO. This is most likely because the AYN is one generation later than the WSS in the breeding process and has already gone through one cycle of selection for wheat stem sawfly resistance. Summary statistics of the location mean, minimum, maximum, standard deviation, heritability and standard error of heritability are also provided (Table 4).

**Figure 5.**
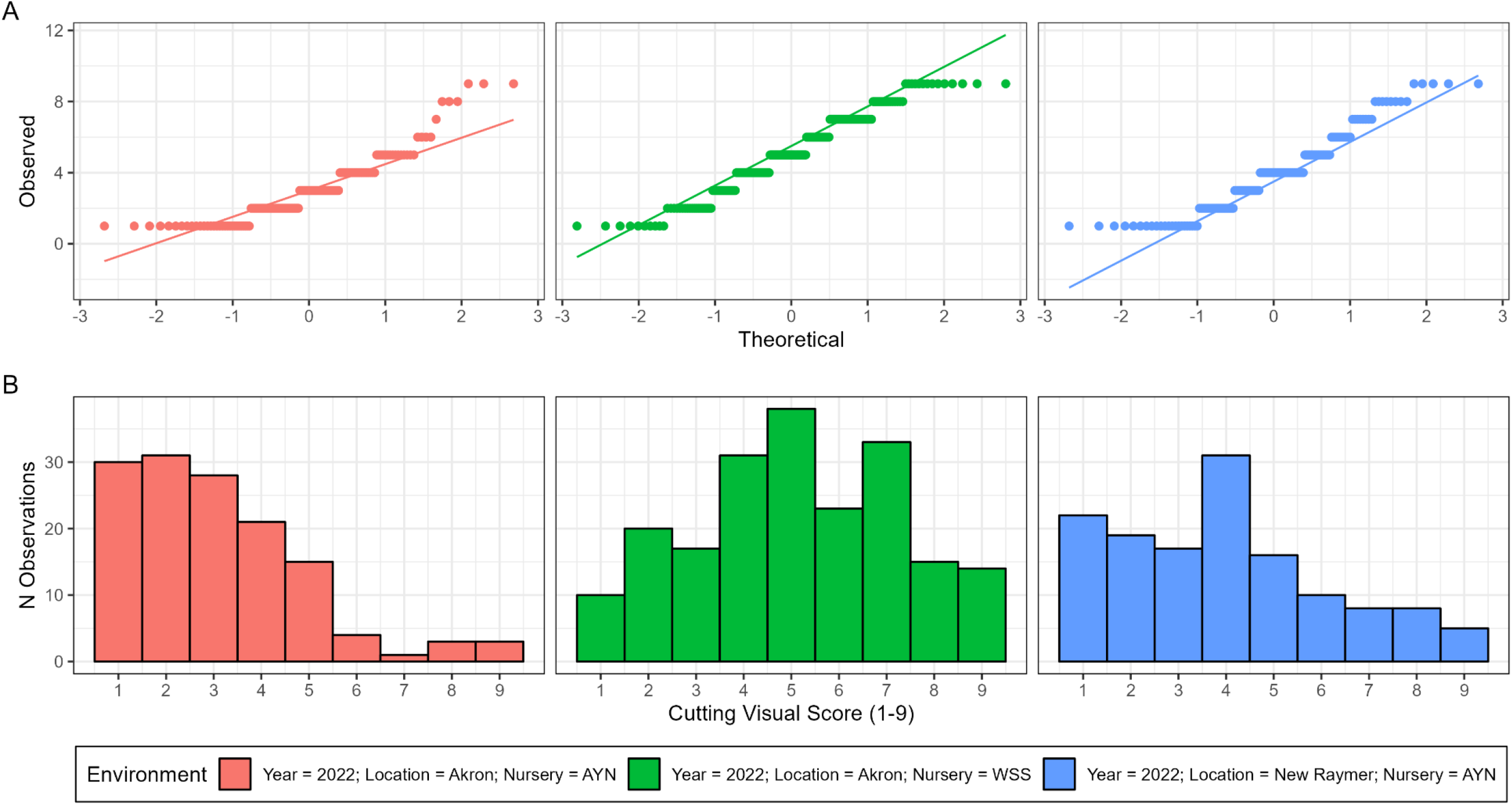
A visualization of (A) qqplots for each locations data and (B) histogram of the cutting visual score within each environment. In panel (A) the y axis represents the observed cutting visual score and the x axis represents the theoretical quantiles. The line going across observation points shows the pattern of expected vs observed visual scores for a normal distribution. Panel (B) displays histograms of each location where the y axis is the count of observations within the bin and the x axis is the cutting visual score. The legend at the bottom of the image displays each environment which corresponds to the color of each subgraph.

**Table 4.** Table of descriptive statistics for each environment.

| Year | Location | Nursery | N Observations | N Genotypes | Min | Mean | Max | SD<sup>a</sup> | $h_g^2$<sup>b</sup> | SE<sup>c</sup> |
| --- | --- | --- | --- | --- | --- | --- | --- | --- | --- | --- |
| 2022 | Akron | AYN | 136 | 107 | 1.00 | 3.07 | 9.00 | 1.86 | 0.62 | 0.10 |
| 2022 | New Raymer | AYN | 136 | 107 | 1.00 | 3.97 | 9.00 | 2.23 | 0.70 | 0.08 |
| 2022 | Akron | WSS | 201 | 185 | 1.00 | 5.12 | 9.00 | 2.18 | 0.49 | 0.11 |

Note: <sup>a</sup> standard deviation of cutting visual score, <sup>b</sup> narrow-sense, per-plot, genomic heritability, <sup>c</sup> standard error of heritability
measurements.

Narrow-sense, per-plot, genomic heritabilities varied among locations. The lowest heritability (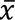 = 0.49 ± 0.11) was observed in Akron, CO for the WSS and the highest (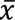 = 0.70 ± 0.08) was observed in New Raymer, CO for the AYN. The mean cutting score in both Akron and New Raymer was lower for the AYN (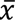 = 3.07 and 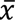 = 3.97, respectively) than Akron for the WSS (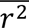 = 5.12), however both nurseries across locations contained visual scores between one and nine. This implies that the generation of selection prior to the AYN did shift the population mean towards resistance, yet it did not cull out all susceptible genotypes, which is expected.

Both predicted and KASP-genotyped *Sst1* haplotype calls had significant effects on cutting score (*P*(*F*) < 0.05). Homozygous *Sst1* and heterozygous *Sst1* individuals did not have substantially different cutting scores when classifying based on either KASP-genotyped or predicted *Sst1* haplotypes. Estimates of *Sst1* effects made by prediction were not significantly different from KASP-genotyped *Sst1* effects within each haplotype (Figure 6). In the case of KASP-genotyped *Sst1* effects, the homozygous wildtype individuals significantly differentiated themselves from both the homozygous *Sst1* and heterozygous individuals; however, predicted haplotypes for homozygous wildtype individuals did not significantly differentiate from heterozygous individuals. This is because the prediction method tended to incorrectly classify homozygous wildtype individuals as heterozygous (Table 3).

**Figure 6.**
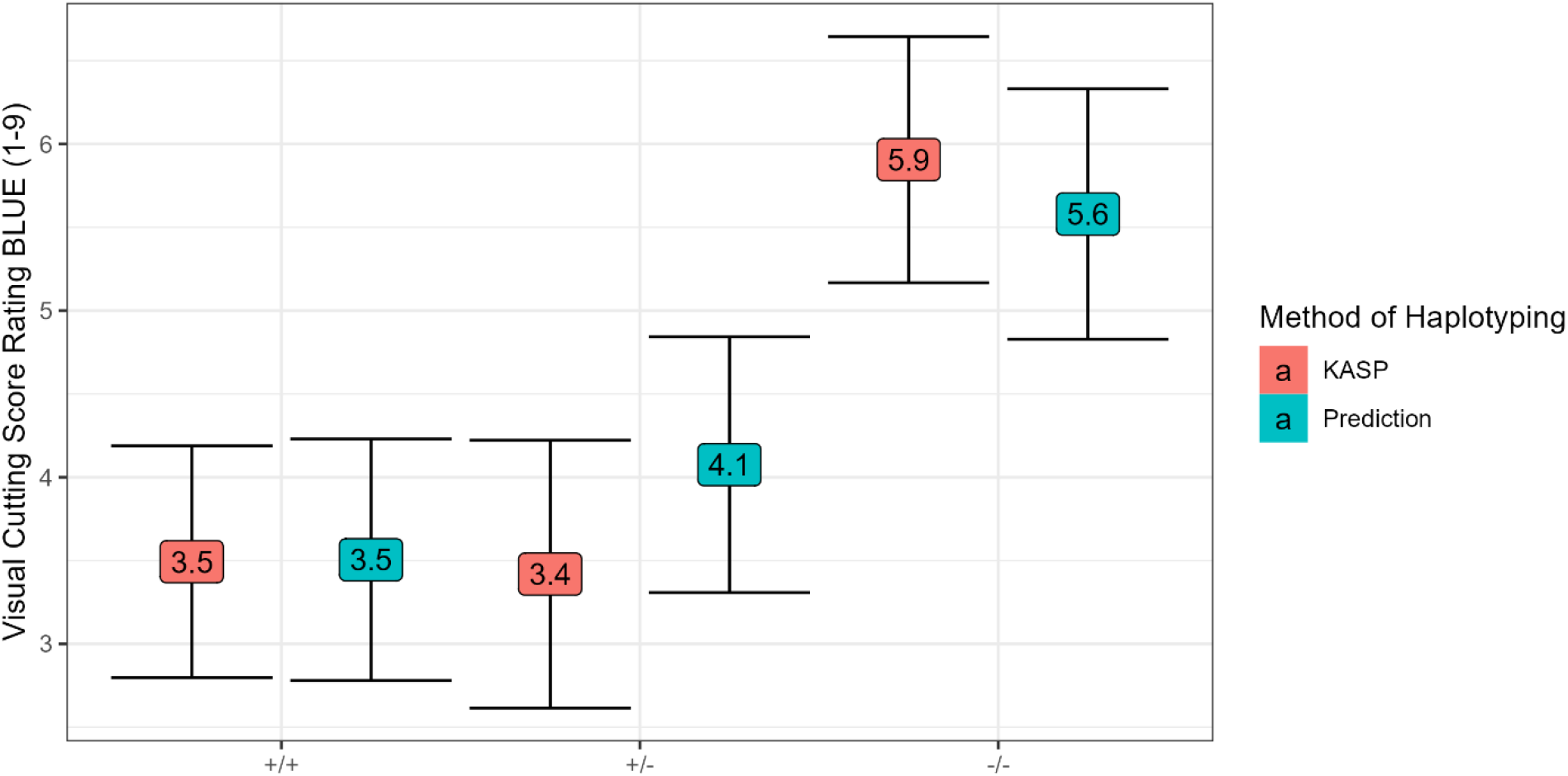
KASP derived *Sst1* haplotype call effects vs predicted *Sst1* haplotype effects. On the x-axis is the allelic state of the *Sst1* locus where +/+ represents individuals homozygous for *Sst1*, +/-represents individuals as heterozygous for the *Sst1* locus, and −/− represents individuals homozygous for the wildtype allele at the *Sst1* locus. The y-axis displays the visual cutting score rating best linear unbiased estimate (BLUE) for the estimated effect of *Sst1*. The legend on the right indicates what color-coded box corresponds to which type of *Sst1* haplotype assignment. The black bars around each point estimate represents a 95% confidence interval about the estimate.

## DISCUSSION

In the current work, we developed an R user accessible R package by the name “HaploCatcher” which can predict haplotypes using historical information derived from molecular marker assays on genome-wide genotyped lines. The function, “auto_locus”, allows users to produce predictions for the many lines submitted for sequencing which are not KASP genotyped on an annual basis. Just as in the work performed by Winn et al (2022), we suggest that this method may be deployed in generations where genome-wide sequencing is performed on a very large number of lines which would otherwise not be screened via PCR based assays for these loci. While these predictions were not perfect in their predictive accuracies (*k* = 1), they were substantial in their predictive ability (*k* ≥ 0.80) and similar in respect to the results of Winn et al (2022) (Landis & Koch, 1977).

Furthermore, this method is directly accessible to breeding programs, researchers, and students due to its development and deployment through R, a free and accessible statistical computing language. Here, we demonstrated that this method is successful in predicting the *Sst1* locus which has a direct impact on improving sawfly resistance in areas threatened by this emerging pest. Moreover, applying this method to resistance loci beyond *Sst1* could lead to further progress in the pyramiding and maintenance of wheat stem sawfly resistance.

The cross-validation results in the current work were similar to those in Winn et al (2022); however, unlike Winn et al (2022) we included the option “include_hets” in the “auto_locus” function, which allows for the prediction of biallelic loci with a heterozygous state. In our results, we observed that both the KNN and RF models were not as capable of identifying heterozygous cases as they were homozygous cases. There may be several reasons for this phenomenon.

Firstly, lines in the CSU program are often initially sequenced in the F_3:5_ generation with recurrent sequencing in each subsequent year. Leaf tissue from ten seeds of each line is bulked and used to prep libraries for sequencing. Therefore, if a heterogenous line for the *Sst1* locus was selected in the F_3_ generation, the DNA extracted may be a small 1:2:1 mixture of *Sst1* haplotypes, and because of this, the sequence of the region may not be truly representative of a heterozygous *Sst1* haplotype, leading to misidentification by this method.

Secondly, we curated KASP data produced by the USDA Central Small Grains genotyping lab over years for training. This data, while highly informative, showed some inconsistency across years. Marker platforms and locus region sizes change over years, and this can lead to unexpected association of markers with the locus. More specifically, we observed that markers 15-20 Mbps away from the known region of the locus were identified as “highly important”. This may be because markers which were used to haplotype the region were not in direct linkage with the causal polymorphism, and this led to the detection of markers on the distal long arm of 3B as important.

Furthermore, we aligned our genetic data to the IWGSC wheat reference genome version 2.0 (Appels et al., 2018). This genome is genetically distant from the wheat germplasm located in the Great Plains area of the United States and may have led to misalignments of sequencing reads, like those potentially observed in the marker importance figure (Figure 3A). Moreover, this genetic data was imputed using Beagle (Browning et al., 2018), which is also not perfectly predictive. Therefore, the summed errors of genomic sequencing method, historical data curation, misalignment, and imputation may have contributed to the lower predictability of heterozygous classes. We therefore suggest that if users can do so, that they produce their own training populations within their own programs and genotype them with a consistent set of markers for best results. Regardless, our proposed models are still substantially informative.

When comparing KASP-based and predicted *Sst1* haplotype call group mean estimates, we observed that the predicted and KASP-based haplotype group means were not significantly different from each other within haplotype. We did observe that homozygous wildtype and heterozygous *Sst1* haplotype group means did not significantly differentiate in the prediction as the RF model used to make this prediction often grouped heterozygous individuals with wildtype haplotypes (Table 3).

Irrespective of these shortcomings, this method provides a way to assess haplotypes of interest, with a measurable margin of error, in generations that would otherwise not be screened for these. More importantly, this package now provides an easily accessible method of pipeline implementation for breeding programs. While targeted sequencing platforms (Lundberg et al., 2013) may reduce the need for a method like this, it will remain useful for programs without access to targeted sequencing platforms or programs missing probes for specific loci of interest. Furthermore, this method can accommodate many different sequencing platforms (diversity arrays, genotyping-by-sequencing, amplicon, etc.), does not require physical position information, and can potentially be widely applied across species. Lastly, we demonstrated that this method can be applied within breeding programs and produce comparable results to PCR based marker calls; more specifically, we showed that this method could be a viable way of screening early development germplasm for the *Sst1* locus, and thus increase the frequency of this locus earlier in the development pipeline.

## CONCLUSIONS

The utility of marker-assisted selection has been vetted through the vast literature available for the method. However, with whole genome sequencing technologies being applied in early generations for use in genomic prediction, there lies an opportunity to acquire data on haplotypes of important loci. The method proposed in Winn et al (2022) allows breeding programs to organize their historical marker-assisted selection data to produce predictive haplotype calls for lines in generations where PCR-based assays for loci of interest are not run due to increased time, labor, and genotyping cost. This can allow breeders to observe locus profiles of potential varieties much earlier in the breeding process than before. Here, we chose wheat stem sawfly – an emerging pest that threatens grower profitability and the dryland cropping agroecosystem– as a test case to demonstrate the effectiveness of this method. We used existing genotypic data sets to deliver breeders precise predictions of the presence of a major resistance gene, *Sst1*, allowing for improved selection for stem sawfly resistance at an earlier generation. With the development of the HaploCatcher package, there is now a freely accessible software for easier implementation of this method in other breeding pipelines.

## ABBREVIATIONS

AYN: Advanced Yield Nursery
BLUE: Best Linear Unbiased Estimate
IWGSC: International Wheat Genome Sequencing Consortium
KASP: Kompetative Allele Specific Polymerase Chain Reaction
KNN: K-Nearest Neighbors
LD: Linkage Disequilibrium
Mbp: Megabase pair
PCR: Polymerase Chain Reaction
RF: Random Forest
RGON: Regional Germplasm Observation Nursery
SNP: Single Nucleotide Polymorphism
SRPN: Southern Regional Performance Nursery
USDA: United States Department of Agriculture
WSS: Wheat Stem Sawfly Solid Stem Panel

## ACKNOWLEDGMENTS

This research was made possible by funds derived from the competitive grant 2022-68013-36439 (WheatCAP) from the USDA National Institute of Food and Agriculture. Mention of trade names or commercial products in this publication is solely for the purpose of providing specific information and does not imply recommendation or endorsement by the US Department of Agriculture. The USDA is an equal opportunity provider and employer.

## DATA AVAILABLILITY

Code and data utilized in this study may be found at <https://github.com/zjwinn/HAPLOCATCHER-A-PACKAGE-FOR-PREDICTION-OF-HAPLOTYPES>. The package developed for this project, “HaploCatcher”, can be directly downloaded to an R installation using devtools::install_github(“zjwinn/HaploCatcher”) to directly install from GitHub or install.packages(“HaploCatcher”) to install from the CRAN database.

